# Toward AI-Resilient Screening of Nucleic Acid Synthesis Orders: Process, Results, and Recommendations

**DOI:** 10.1101/2024.12.02.626439

**Authors:** Bruce J. Wittmann, Tessa Alexanian, Craig Bartling, Jacob Beal, Adam Clore, James Diggans, Kevin Flyangolts, Bryan T. Gemler, Tom Mitchell, Steven T. Murphy, Nicole E. Wheeler, Eric Horvitz

## Abstract

Fast-moving advances in AI-assisted protein engineering are enabling breakthroughs in the life sciences that promise numerous beneficial applications. At the same time, these new capabilities are creating potential biosecurity challenges by providing new pathways to intentional or accidental synthesis of genes that encode hazardous proteins. The synthesis of nucleic acids is a key choke point in the AI-assisted protein engineering pipeline as it is where digital designs are transformed into physical instructions that can produce potentially harmful proteins. Thus, one focus for efforts to enhance biosecurity in the face of new AI-enabled capabilities is on bolstering the screening of orders by nucleic acid synthesis providers. We describe a multistakeholder, cross-sector effort to address biosecurity challenges with uses of AI-powered biological design tools to reformulate naturally occurring proteins of concern to create *synthetic homologs* that have low sequence identity to the wild-type proteins. We evaluated the abilities of traditional nucleic acid biosecurity screening tools to detect these synthetic homologs and found that, of tools tested, not all could previously detect such AI-redesigned sequences reliably. However, as we report, patches were built and deployed to improve detection rates over the course of the project, resulting in a final mean detection rate over tools of 97% of the synthetic homologs that were determined, using in-silico metrics, to be more likely to retain wild-type-like function. Finally, we make recommendations on approaches for studying and addressing the rising risk of adversarial AI-assisted protein engineering attacks like the one we identified and worked to mitigate.

## Introduction

Recent advances in artificial intelligence (AI) have enabled a series of astonishing breakthroughs in the life sciences. One of the most promising areas, with early results and expectations for more on the horizon, is the application of machine learning to the understanding and design of proteins,^1,2^ with a growing array of AI tools being harnessed to modify known proteins and design novel proteins. These tools help scientists to cut through combinatorial design spaces to reach goals of specific structures and functions.^3–9^ They have already been used to create new vaccines, antibiotics, cancer therapeutics, and insights into the machinery of life and, indeed, it seems that we are just at the beginning of a new era in biology and medicine.^1,2,5,9–17^ There is great promise ahead for harnessing AI-powered biological design to cure diseases and to nurture wellness, vibrancy, and longevity.

At the same time, however, like other technical advances, the powers of AI are dual use: Although they hold promise for valuable innovations, AI tools for protein design also carry the risk of misuse—whether by malicious actors with intent, or by careless or inexperienced users inadvertently causing harm.^18,19^ So, as we move forward with AI-powered biological design research and development, we must work in parallel to identify and address biosecurity implications of these advances.

Concerns about bad actors co-opting innovations in biology for nefarious means are not new, and a number of safeguards have been implemented to minimize risks.^20–22^ For example, screening efforts have been employed at the juncture between digital designs for proteins and procurement of the nucleic acid sequences required to generate them in the laboratory; many companies that provide researchers with synthetic nucleic acids employ biosecurity screening software (BSS) to identify customer orders containing sequences that may be subject to governmental regulation or that may otherwise pose concerns about potential misuse.^21^ Such screening aims to prevent the construction and proliferation of genes that encode proteins of concern.

To date, BSS systems used by nucleic acid synthesis companies have largely relied on identifying sequence similarities between ordered sequences and sequences unique to regulated toxins, viruses, or other organisms. Such an approach is specified in guidance from the U.S. Department of Health and Human Services (HHS), which was first published in 2010 and updated in October of 2023.^23,24^ With the ever-increasing sophistication and accessibility of generative AI in the biosciences, however, questions are raised regarding the robustness of *identity-based* approaches to the new capabilities that AI enables.^18,25–30^

Review and updating of nucleic acid screening methods in light of AI advances was called for as part of the U.S. President’s Executive Order on the Safe, Secure, and Trustworthy Development and Use of Artificial Intelligence in October 2023.^24,31,32^ Following the issuance of the executive order, the White House Office of Science and Technology Policy (OSTP) assembled representatives from multiple agencies to develop a response to the call. In April 2024, OSTP released the Framework on Nucleic Acid Synthesis Screening based upon the updated HHS guidance, including an updated window size for identifying subsequences of concern. The revised requirements continue to focus on sequence identity. However, the guidance asks for ongoing review and updating of the framework via the creation of a U.S. interagency group charged with assessing the scientific state of the art before October 13, 2026, and recommending any updates as necessary.

The multistakeholder project reported here (overview in Figure 1), explores mitigation of potential biosecurity risks associated with rising challenges to traditional nucleic acid screening based on sequence identity. We investigated a potential vulnerability of identity-based screening of sequences in the face of AI-assisted protein design (AIPD) toolchains that facilitate the diversification of a protein’s amino acid sequence while seeking to maintain its function.^4,6,8,9,33,34^ We found in a confidential process that several BSS, including screening methods in use at major nucleic acid suppliers, could not reliably detect such AI-reformulated toxins and viral proteins. However, we showed that the screening machinery could be patched using insights from this effort to improve their ability to detect AI-designed variants—or *synthetic homologs*—of proteins of concern.

**Figure 1.**
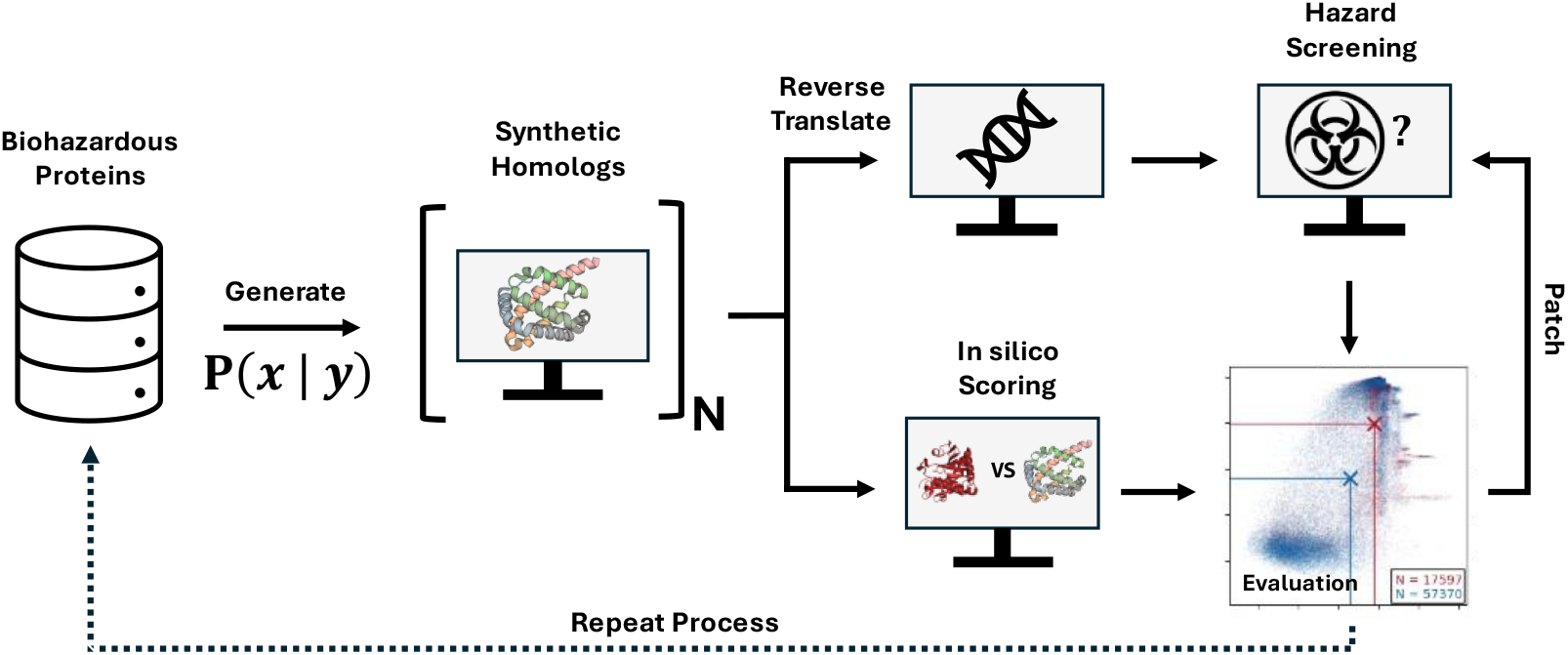
Summary of AIPD red-teaming workflow. Following a framing study that demonstrated key methods and concerns, a list of biohazardous proteins of concern spanning various toxins and pathogen virulence factors was compiled. Next, these proteins were used as templates to generate synthetic homologs in silico. The workflow was then split into two independent paths: In the first (upper), the reformulated protein sequences were reverse-translated into DNA before being evaluated by the biosecurity screening software (BSS) of participating providers. Two reverse translation strategies were used in this work, one that employed a simple random mapping between amino acid identity and codon, and another that further obscured the identity of the reformulated protein. In the second (lower) path, the structures of all generated and wild-type proteins were predicted. Using these predicted structures, the in-silico metrics, ΔpLDDT and TM-Score, were calculated; these metrics can be thought of as providing a measure of the probability that a synthetic homolog will maintain the structure of its wild-type template, which in theory is correlated with maintaining function. The two paths were then converged, and the results of the various BSS were mapped to the in-silico metrics. These results were used to evaluate BSS performance and to update (patch) the BSS to address identified weaknesses. As protein design models improve, we anticipate that this process and others like it will need to be repeated to ensure that BSS tools remain effective.

Beyond doing a technical deep dive into the opportunity to bolster the resiliency of BSS to advances in AIPD, we have explored identifying valuable processes for AI biosecurity red teaming, response, and mitigation. We share our approach, which was informed by cybersecurity practices for addressing detected vulnerabilities, in the hope that the phases of study, testing, and response can serve as an exemplar for future work on other AI-centric biosecurity concerns. Our process included securely pursuing, identifying, and characterizing a biosecurity challenge enabled by AIPD, building a trusted multiparty-stakeholder team, maintaining confidentiality over an extended period of time on details of discovered concerns while study of the challenge and its mitigations progressed, alerting relevant government agencies about the challenge and on progress of mitigations, and sharing details on the challenge and best practices on mitigation with a broad group of screening providers in advance of public disclosure once findings and approaches had become clear. Finally, we deliberated about the value versus potential hazards of publicly sharing versus withholding information about the project, including questions about if, when, which, and with whom details regarding our methods, findings, and process should be shared, up to and including our decision to release this publication.

## Considerations and Process

In advance of presenting details on a detected AI-enabled biosecurity vulnerability, and methods, results, and mitigations developed, we share background on the process that we undertook. This includes activities in advance of performing the study and deliberations at its end, in advance of releasing information about our methods and results.

Our study centers on a careful AI red-teaming analysis performed with the awareness of leads at relevant U.S. government agencies and in the context of the opportunities and challenges with AI and biosecurity called out in the U.S. Executive Order on AI. The research was performed in the context of voluntary commitments made to the White House on AI safety and security, announced earlier by several organizations engaged in leading-edge AI research and development.

We note that, per the high-stakes and potentially costly outcomes of the research, AI and biosecurity red-teaming efforts need to be compliant with existing international and national agreements. The studies must aim to promote peaceful uses of technology with a goal of preserving life and health. As our study progressed, we stayed in contact with leads engaged on AI and biosecurity at key U.S. government agencies.

We took inspiration from processes established by the cybersecurity community for addressing zero-day situations, referred to as a *computer emergency and response team* (CERT) methodology.^35,36^ We withheld publication of an identified AI-enabled biosecurity vulnerability until a “patch” could be developed and distributed, engaged with key stakeholders, built an expanded team to develop a deeper characterization and understanding of BSS vulnerability, and worked to mitigate the weaknesses. Following the development of a solution, we disclosed the vulnerability, along with information about mitigations in a confidential process.

Finally, we conducted a multistakeholder deliberation with biosecurity experts across sectors to discuss the scientific and policy benefits versus the potential information risks of publicly sharing the methods, results, and overall process of the study. Those discussions guided decisions about the content of this manuscript, including details withheld. Overall guidance received was that public sharing of top-level information on the study, including a summary of methods, results, and overall process, would have strongly positive benefits, per advancing understandings and future valuable efforts by the scientific community, civil society, and government agencies on biosecurity amidst rising powers of AI technologies. To minimize risk, we decided to keep confidential a constellation of details, including the AI tools used to generate synthetic homologs, the identity of the set of natural proteins considered in our diversification studies, and details of the generative and analytical methods employed over the course of the study.

## Results

We shall now review results from an initial framing study, followed by a larger study sparked by the initial analysis.

### Framing Study

Our project was stimulated by an initial end-to-end adversarial AIPD framing study performed by B.J.W. and E.H. in October 2023. The goal of the framing study was to explore AI design capabilities that could provide an example for focusing discussions at a forthcoming summit of protein engineering scientists on biosecurity at the University of Washington later that month.^37^ In the framing study, widely available AIPD tools were used to demonstrate in silico how a protein of concern could be reformulated into thousands of synthetic homologs with alternate sequences. The sequences resulting from this reformulation process were not tested in vitro for preserved function; however, literature evidence suggests that at least a subset of the diversified analogs created could retain some or all function were they to be physically synthesized, and that some of the new proteins could even have increased function relative to the wild type.^3,4,6,7,9,33,34,38,39^ Regardless, even if no designs retained function, given the rapid pace of advancement of AIPD tools over the last five years, it is not difficult to imagine a future where this is not the case and most generated proteins function. Addressing biosecurity risks now—whether they be anticipated or salient—is vital to ensuring that the benefits of AIPD are realized while minimizing the risk of harm.

The leads of the framing study reached out to two prominent nucleic acid synthesis companies and confirmed that BSS in use at their organizations could not reliably detect reverse-translated DNA sequences encoding the synthetic homologs. However, it was also shown that these organizations’ screening software could be updated with a “patch” to make the screening more resilient to AI-reformulated sequences, leveraging insights garnered from the set of synthetic homologs.

The details on the framing study were described in a whitepaper that was shared with several trusted colleagues in the protein design community who attended the biosecurity meeting at the University of Washington. The leads of the framing study also informed the chair of the International Gene Synthesis Consortium (IGSC), an industry-led group of gene synthesis companies and organizations formed to design and apply a common protocol to screen both the sequences of synthetic gene orders and the customers who place them.^21^ Further, U.S. government biosecurity leads were informed about the potential vulnerability identified by the study and the planned follow-on studies.

Despite the lack of wet-lab studies confirming functionality of the synthetic homologs produced in the framing study, we believe that the dramatic—though perhaps not necessarily unexpected—failure of the tested BSS tools represented a type of biological *zero day*, a term commonly used in the cybersecurity domain to describe software or hardware vulnerabilities that were previously undescribed and for which no patch exists at the time of discovery.^40^ As noted earlier, we moved forward with a modified CERT process in the spirit of the approach taken within the cybersecurity community for handling a cybersecurity zero day. As part of this process, we made a decision to keep the whitepaper describing the framing study confidential (beyond the limited sharing with appropriate parties as described above) and to pursue a more intensive effort to further characterize and develop countermeasures to the biosecurity risk of using AI to generate synthetic homologs of biohazardous proteins.

In the larger, follow-on study, the results of which are presented in the remainder of this paper, the circle of collaborating organizations was increased to include additional DNA synthesis companies and biosecurity screening tool providers, the identities of which are represented by the home institutions of authors of this paper. We also expanded the single protein analysis via a consensus process to a much longer list of proteins of concern. Using these proteins of concern as templates, we generated synthetic homologs in silico to evaluate the ability of BSS systems used by major nucleic acid synthesis and screening companies to identify those new proteins as potential hazards. We confirmed that several BSS strategies in use at the time of the framing study could not identify AI-reformulated variants of proteins of concern, with up to 100% of variants from certain proteins passing undetected through at least one BSS. As we conclude this larger phase of study, however, we confirm that patched BSS are now far more capable of identifying such synthetic homologs, detecting 97% of proteins deemed to have the potential of maintaining function according to in-silico metrics. Capture rates are significantly higher with orders containing multiple sequences, as might be expected with attempts to generate functional synthetic homologs, given the recognized challenges of achieving functionality even when structure is largely maintained.

In keeping with established cybersecurity practices, we informed organizations that deploy BSS in advance of making this larger study and its results publicly available, providing recommended screening software updates and pointers to best practices identified as part of this work to members of the IGSC on July 30, 2024. Now that the broader project has been completed, we additionally make the initial framing paper from October 2023 available as the appendix to this manuscript.^41^

### In-Silico Protein Reformulation Pipeline

Most BSS systems have operated by searching a sequence for fragments of controlled nucleic acid or protein sequences. If a fragment is a better match to a sequence of concern than to any sequence from other, non-controlled organisms, then the sequence is flagged as potentially “of concern” and queued for expert review. This process, referred to as *best match*, was originally defined in the since-updated sequence screening guidelines published by HHS in 2010.^23^

BSS systems in use today are the result of refinement over time to improve resilience to newly emerging exploits. For example, at one point, it was challenging to change the wild-type nucleic acid sequence encoding a protein while maintaining protein expression, but the emergence of *codon optimization* and *codon reassignment* tools has allowed for the design of nucleic acid sequences that have low identity to controlled nucleic acid sequences while encoding an identical amino acid sequence.^42^ Consequently, current BSS tools have been made “codon resilient” by adding six-frame translation of nucleic acid sequences and checking for sequence similarity to amino acid sequences of concern.

AIPD tools expose vulnerabilities in BSS in a similar manner to codon exploits. However, rather than changing the nucleic acid sequence while maintaining the amino acid sequence, the AI methods—specifically, those using protein sequence generative models (PSGMs)—enable bad actors or unsuspecting researchers to change the amino acid sequence while pursuing maintenance of function. In doing so, the resultant proteins can have limited amino acid sequence identity to a controlled amino acid sequence, and so can be difficult to detect by “best match” sequence similarity. Indeed, AIPD is now making it possible to generate function-equivalent, sequence-diverse proteins potentially hundreds of mutations away from the closest known natural protein.^4,33,34,39^

To investigate the level of resilience of BSS to adversarial AIPD attacks, we developed a red teaming workflow for in-silico generation and evaluation of synthetic homologs of proteins of concern (Figure 1). Importantly, to best represent in our biosecurity red-teaming effort the tools available to those who would use AIPD for harm,^18^ we relied solely on publicly available, open-source PSGMs, making no modifications to the underlying AI models that power them. The results from using three of these PSGMs in our red teaming workflow are provided in the rest of this work.

We note that none of the PSGMs used in our study are foolproof; there is much to learn about why some generated protein designs fail while others have the targeted function. Indeed, depending on the protein being reformulated, we expect varying degrees of success regarding the goal of maintaining function. For instance, catalytic activity relies on more precise positioning of atoms within an enzyme’s active site than does binding activity on the positioning of atoms at a binding interface. Thus, catalytic activity will likely be more fragile in the face of high mutational loads than binding activity. Similarly, it is not uncommon for one or a few residues in a protein to be critical for its function; and because, by definition, the PSGMs used here incorporate a degree of randomness when generating a new sequence, it is possible that these are not maintained in generated variants. Thus, the more essential residues there are in a protein, the less likely it is that reformulated variants will maintain wild-type-like function.

To account for the fact that different proteins have different sensitivities to mutational load, we configured our PSGMs such that each could be provided with a set of fixed amino acids at pre-specified positions. When run in this “restricted” mode, the models produce sequences that are less sequence diverse but more likely to function. This also means that sequences generated in restricted mode would likely be easier to detect using traditional BSS. Thus, with currently available AIPD tools, there is a tradeoff between the probability that a generated protein maintains function and the ease with which that protein could be detected by BSS.

As noted earlier, we did not construct and test any of the variants of proteins generated as part of this study in a laboratory setting, given ethical concerns, international norms, and the Biological Weapons Convention that restricts the construction of materials that could result in the weaponization of biology.^43^ Such confirmatory work and real-world calibration via lab-based studies, if conducted, must be carried out with appropriate security, transparency, and oversight. Instead, we relied on in-silico metrics in this study. Specifically, we calculated a predicted structure and structure-level predicted local distance difference test (pLDDT, which is a measure of prediction confidence) for every wild-type template protein and in silico-generated synthetic homolog. Then, we used those calculations to calculate both a template modeling score (TM-Score, which is a structural similarity metric) and ΔpLDDT for each synthetic homolog (Methods: In Silico Evaluation of Generated Proteins).

A synthetic homolog predicted to maintain the folding structure of its wild-type template is more likely to maintain function. Thus, synthetic homologs with a high TM-Score are more likely to maintain function. Similarly, we expect protein structure prediction models generally to report a higher pLDDT for sequences similar to those to which it was exposed during training. This is because pLDDT is a confidence metric, and models tend to be more confident when making predictions on inputs that are in-distribution with regard to their training data. As the model used for predicting protein structures was trained using proteins known to have some function, pLDDT can be thought of as a measure of how “realistic” an input sequence is, conditioned on the known space of possible proteins. In other words, a synthetic homolog with a higher pLDDT (and thus a higher ΔpLDDT) is more similar to known, functional proteins and so is more likely to maintain the original function. Indeed, previous studies have used pLDDT and structural similarity metrics like TM-Score to identify successfully reformulated (benign) proteins.^7,9,33,38^

We stress that a high ΔpLDDT and high TM-Score do not guarantee that a reformulated variant will maintain function. Protein structure predictions are themselves imperfect and, even given ground-truth structures, a high structural similarity between a variant and its template is merely necessary, not sufficient, for that variant to maintain function. Literature evidence, personal communications with other experts in AIPD, as well as unpublished, analogous, in vitro-tested follow-on work to this study using benign proteins as proxies for hazardous proteins suggest that synthetic homologs with low ΔpLDDT and TM-Scores are extremely unlikely to retain function; the rate of function retention in synthetic homologs with high ΔpLDDT and TM-Scores, in contrast, varies depending on the template protein, as expected based on the discussion above. Throughout this work, then, variants with higher ΔpLDDT and TM-Scores should only be viewed as *more likely* to maintain function than those with lower values, and only when comparing variants derived from the same template protein. While this is a limitation of our analysis, we stress that the goal of this effort was to improve the ability of BSS to detect *potential* sequences of concern, not necessarily to detect confirmed *functional* sequences of concern. With this risk-reduction goal in mind, we believe that in-silico metrics are sufficient for evaluating the relative performance of BSS implementations.

### In-Silico Reformulation

As part of our red teaming workflow, we generated tens of thousands of synthetic homologs in silico from the aforementioned list of proteins of concern. (Methods: Variant Generation). Figure 2 shows the distributions of the TM-Score, ΔpLDDT, and sequence-identity-to-the-template for the synthetic homologs generated from three PSGMs. In general, we see that Model 3 produces synthetic homologs that have the lowest sequence identity relative to their wild-type templates and, according to our in-silico metrics, are the least likely to maintain the function of their wild-type templates. The variants generated by Model 2 have the next-lowest sequence identities and inferred probabilities of retaining function, followed in turn by those generated by Model 1.

**Figure 2.**
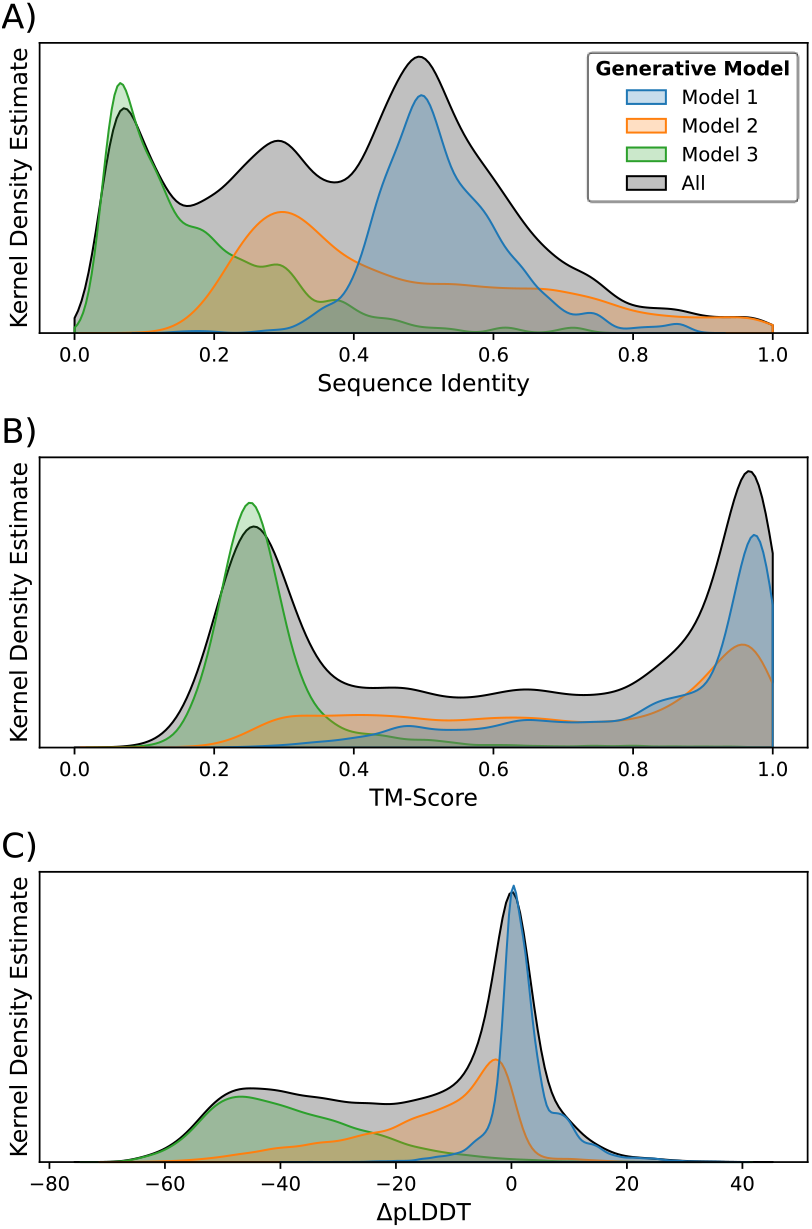
Distribution of metrics for the synthetic homologs generated in silico by three protein sequence generative models. (A) Distribution of sequence identities between the wild-type proteins and the derived reformulated variants. (B) Distribution of TM-Scores between the wild-type proteins and derived reformulated variants. (C) Distribution of the difference in pLDDT between the wild-type proteins and the derived reformulated variants. In general, we see that Model 3 generates proteins that are least likely to function and with lowest sequence identity compared to the wild-type templates, followed by Model 2, and then by Model 1.

### Ability to Detect AI-Reformulated Proteins Varied by Screening Tool

To synthesize any biohazardous protein, a bad actor would first need to order a gene (or fragment thereof) encoding that protein from one of numerous synthetic nucleic acid providers, then express that nucleic acid in a host organism or in-vitro system to produce the protein. Thus, the protein sequences described in the previous section were reverse translated into DNA before we used them to evaluate participating providers’ BSS. Once reverse translation was complete, the resulting DNA sequences were sent to the participating BSS developers, who ran the sequences through their software (Methods: Screening DNA for Potentially Hazardous Proteins).

The results of runs performed using BSS that were used in production at the time that the framing study was performed in October 2023 can be seen in the first column of Figure 3, where we visualize the results in terms of the TM-Score and ΔpLDDT of the synthetic homologs. A synthetic homolog with both a high TM-Score and high ΔpLDDT is more likely to maintain the function of its wild-type template. Thus, we consider it important for BSS to flag most of the sequences in the top right of the subplots of the first and second columns of Figure 3 (i.e., the top right of the subplots should be red). It is important to note that, when applied to standard nucleic acid synthesis orders, most of which will not contain potentially hazardous sequences, the false positive rate must be kept low to avoid unnecessary delays in order fulfillment and minimize the economic burden imposed by screening on the nucleic acid synthesis industry. Thus, we do not necessarily want to see BSS flagging proteins in the lower-left corner of these plots. Still, given the potential for the proteins studied in this work to cause harm as well as the uncertainty inherent to our in-silico metrics, it is more important here to see the upper right colored mostly red than the lower left colored mostly blue. To be most confident in a BSS system, we should *not* see its performance characterized by blue patches in the upper right of the first or second columns of Figure 3.

**Figure 3.**
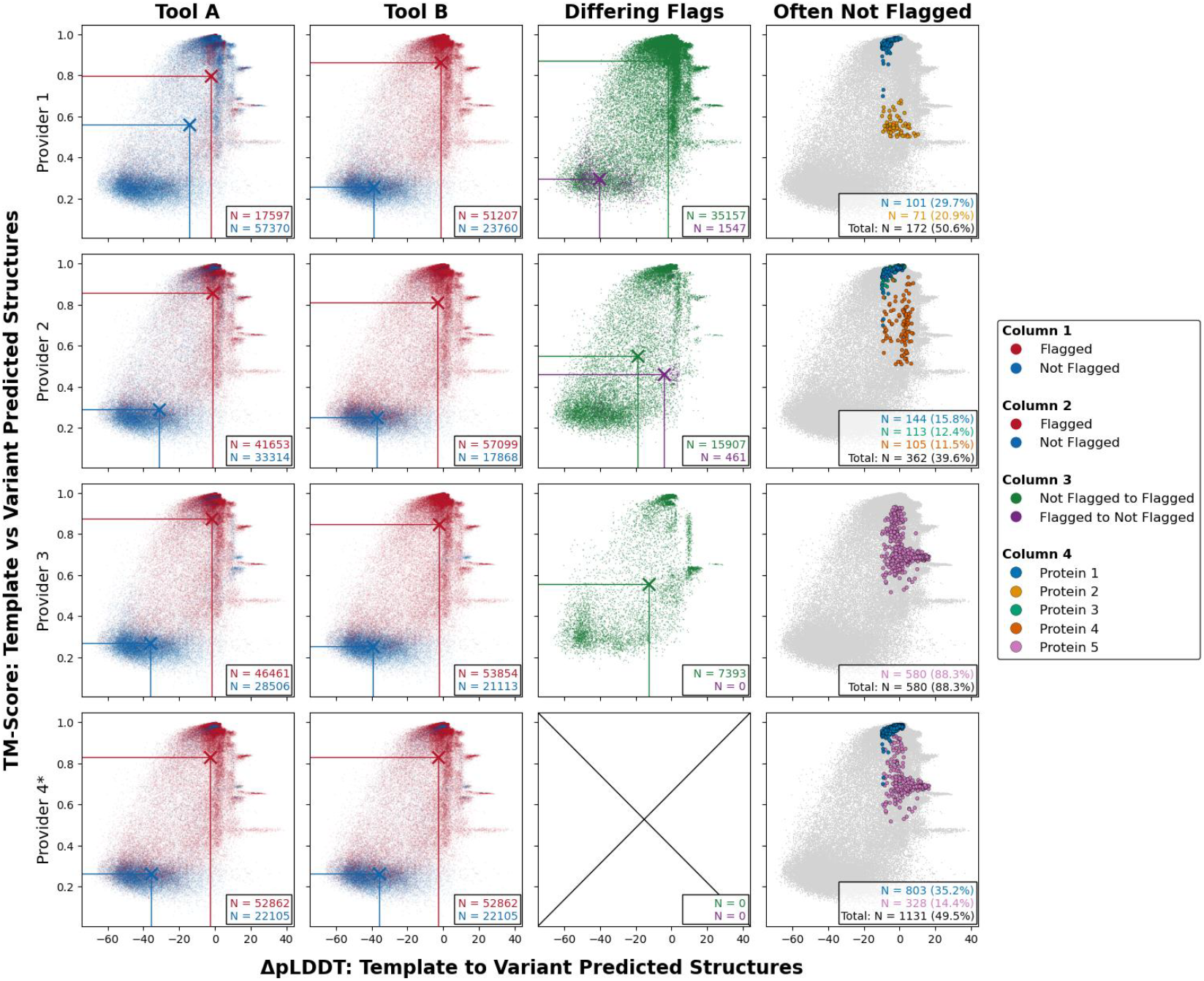
In-silico metrics and flags for all proteins, tools, and participating providers. The x-axes give the ΔpLDDT and the y-axes give the TM-Score. Each row corresponds to a different participating biosecurity screening software (BSS) provider. Column 1 (Tool A): Results from BSS strategies available when the framing study was performed in October 2023, with red dots indicating a protein that was flagged as hazardous and blue dots indicating one that was not. X marks indicate the median values for the in-silico metrics. Column 2 (Tool B): Results of BSS updated to be more capable of identifying AI-reformulated protein sequences over the course of the study. Results from Column 1 are copied to Column 2 for Provider 4 as no updates were made to this tool between the date of the framing study and the writing of this report. Column 3 (Differing Flags): Changes in flags between Columns 1 and 2. Green dots indicate proteins that were not flagged in Column 1 but that are in Column 2, and purple dots indicate the opposite. Proteins whose flags did not change are not plotted in this column. X marks again indicate the median values. Column 4 (Often Not Flagged): Frequently unflagged proteins with high in-silico metrics (TM-Score > 0.5 and ΔpLDDT > -10) for tools whose results are presented in Column 2. Boxes in the lower right corner of plots in this column give the number of these proteins and the percentage of all unflagged high-in-silico-metric “misses” that this number constitutes. Only proteins constituting >10% of misses were highlighted in this column. See also, Table 1.

Overall, the results displayed in Figure 3, Column 1 were mixed, with all BSS tools not flagging some proteins that had high in-silico metrics, but the total number of such unflagged high-scoring proteins varying from tool to tool. This was not the case when the BSS were applied to the wild-type templates used to build the reformulated variants, however, where we observed near-perfect sensitivity and specificity for all tools (Table 1). Together, these results indicate that, while all tested BSS worked well for wild-type proteins, given the newfound capabilities of PSGMs, not all were sufficient for identifying DNA synthesis orders that contained genes encoding potentially harmful proteins reformulated using AIPD strategies.

**Table 1.** Confusion matrices for each tested biosecurity screening software (BSS) tool against the synthetic homologs and control sequences. A “+” in a tool row indicates the frequency with which a group of sequences was flagged as potentially hazardous, while a “-” indicates the frequency with which a group of sequences was not flagged. A “+” in the “More Probably Functional?” column indicates that the reported frequency was calculated against the group of sequences with high in-silico metrics (TM-Score > 0.5 and ΔpLDDT > -10) while a “-” indicates the opposite. A “+” in the “Control Type” column indicates that the reported frequency was calculated against the wild-type sequences used as templates to build synthetic homologs. A “-” in the “Control Type” column indicates that the reported frequency was calculated against reformulated sequences that were generated from PSGMs conditioned on no other information than the length of a protein of concern’s sequence. Overall, we see that all BSS that were updated were improved in their ability to detect more-probably-functional sequences. We also see that these updates did not negatively impact the tools’ abilities to distinguish between known controlled sequences (the positive controls) and presumed benign sequences (the negative controls).

|  |  |  | More Probably Functional? |  | Control Type |  |
| --- | --- | --- | --- | --- | --- | --- |
|  |  |  | - | + | - | + |
| Label Predicted by BSS | Tool 1A | - | 0.8335 | 0.6806 | 0.9976 | 0.0185 |
|  |  | + | 0.1665 | 0.3194 | 0.0024 | 0.9815 |
|  | Tool 1B | - | 0.5739 | 0.0099 | 0.9780 | 0.0000 |
|  |  | + | 0.4261 | 0.9901 | 0.0220 | 1.0000 |
|  | Tool 2A | - | 0.6654 | 0.1797 | 0.9998 | 0.0494 |
|  |  | + | 0.3346 | 0.8203 | 0.0002 | 0.9506 |
|  | Tool 2B | - | 0.4155 | 0.0266 | 0.9991 | 0.0000 |
|  |  | + | 0.5845 | 0.9734 | 0.0009 | 1.0000 |
|  | Tool 3A | - | 0.6084 | 0.1075 | 0.9953 | 0.0556 |
|  |  | + | 0.3916 | 0.8925 | 0.0047 | 0.9444 |
|  | Tool 3B | - | 0.5013 | 0.0192 | 0.9953 | 0.0123 |
|  |  | + | 0.4987 | 0.9808 | 0.0047 | 0.9877 |
|  | Tool 4 | - | 0.4857 | 0.0666 | 0.9809 | 0.0247 |
|  |  | + | 0.5143 | 0.9334 | 0.0191 | 0.9753 |

Column 2, Rows 1–3 of Figure 3 show the results of BSS updated either in response to the results presented in Column 1 or in response to the results reported in the framing study. Provider 4, whose results are presented in Row 4, did not perform updates to their tool for reasons discussed later, so its results from Column 1 were copied into Column 2; the overall strategy used by this provider at the time of the framing study was identical to the strategy used at the time of writing. After making changes to their BSS strategies, Providers 1–3 became more capable of identifying AI-reformulated variants and, in some cases, also at identifying the wild-type templates (Table 1).

The types of changes made to the BSS to improve their resilience to reformulated variants are reflected in the distributions of changes to labels (Figure 3, Column 3). For instance, Tools 1A and 1B rely on completely different strategies to one another; Tool 1A can be considered a traditional BSS as it relies on sequence identity matches to a list of proteins of concern to determine whether a sequence should be flagged as hazardous, whereas Tool 1B augments that approach with a model designed to recognize signatures of AI-generated variants of proteins of concern. Looking at the Figure 3, Column 3 results for the latter tool, we see that most variants that go from unflagged to flagged reside in the upper right of the plot, where we expect synthetic homologs to have both sequence and structure signatures more similar to their associated wild-type templates (corresponding to higher ΔpLDDT and TM-Score, respectively, as discussed earlier), and most variants that go from flagged to unflagged reside in the lower left of the plot, where we expect synthetic homologs to have less similar signatures to their templates. Conversely, Tools 2A and 2B as well as 3A and 3B rely on similar mechanisms to one another, but with parameters updated between the A-series and B-series to make Tools 2B and 3B more sensitive to low-sequence-identity matches to sequences of concern. In this case, the proteins moving from unflagged to flagged and vice versa are more evenly distributed across the in-silico metric space, corresponding to a change in the thresholds deemed necessary for assigning each label.

### Screening Software Flags 97% of More-Probably-Functional Reformulated Variants

None of the existing or updated BSS could identify all proteins with high in-silico measures. If we define a “more-probably-functional” protein as one with a TM-Score above 0.5 (a value typically cited as a threshold for whether two proteins are likely to share the same overall fold^44^) and a ΔpLDDT above -10 (a more arbitrarily chosen cutoff that, when combined with the TM-Score cutoff of 0.5, isolates the cluster of proteins in the top right of the subplots of Figure 3, Columns 1 – 2), then Tool 1B does not flag 340 (∼1%) of these, Tool 2B does not flag 913 (∼3%), Tool 3B 659 (∼2%), and Tool 4 2287 (∼7%), corresponding to an average of ∼3% over tools (or, conversely, an average flag rate of 97% over tools). We again stress that we should not expect all (nor even, necessarily, a majority) of these more-probably-functional proteins to actually be functional. The cutoffs were selected to further simulate potential behavior of an adversarial actor, who would, as is common practice when selecting generated protein sequences for analysis in vitro, likely employ an in-silico pre-screen to filter candidates. We also selected more generous cutoff values than those typically used in pre-screens, given the potential for a truly functional reformulated variant to cause harm. However, even if all more-probably-functional variants were in fact functional, the rates at which they were not flagged by the tools are comparable to the false negative rates of the screening tools against the wild-type proteins of concern (Table 1). In other words, the BSS strategies in use as of now by the providers participating in this study (and assumedly other BBS providers, who have by now been alerted confidentially to the potential synthetic homolog vulnerability), are at least as effective at identifying AI-reformulated hazardous sequences as they are at identifying wild-type sequences of concern.

It is still important to analyze why some more-probably-functional sequences were not flagged (referred to as *missed* sequences from hereon). At first, and as discussed earlier, we might assume that these missed sequences belong to reformulated variants with a lower sequence identity to their template. Interestingly, however, while the missed variants did indeed tend to have lower sequence identity relative to their template sequence, this was not the case for all of them. We can see this in Figure 4, which plots the distributions of amino acid sequence identity to the template for all more-probably-functional variants grouped by whether those variants were flagged. Specifically, this figure shows that all BSS did not flag a sizable number of sequences with high amino acid sequence identity to template, where “high” is defined as >60% and was chosen to capture the tail of the distribution of sequence identities (Figure 2A). Indeed, for Tool 3B, more-probably-functional sequences that were not flagged tended to have a *higher* sequence identity to their template than did the flagged ones. While perhaps initially surprising, if we take a closer look at the missed sequences, the reason for the pattern becomes clear: As displayed in the last column of Figure 3, the majority of the missed variants are derived from just a few template proteins, an effect that tends to become more pronounced at higher sequence identities.

**Figure 4.**
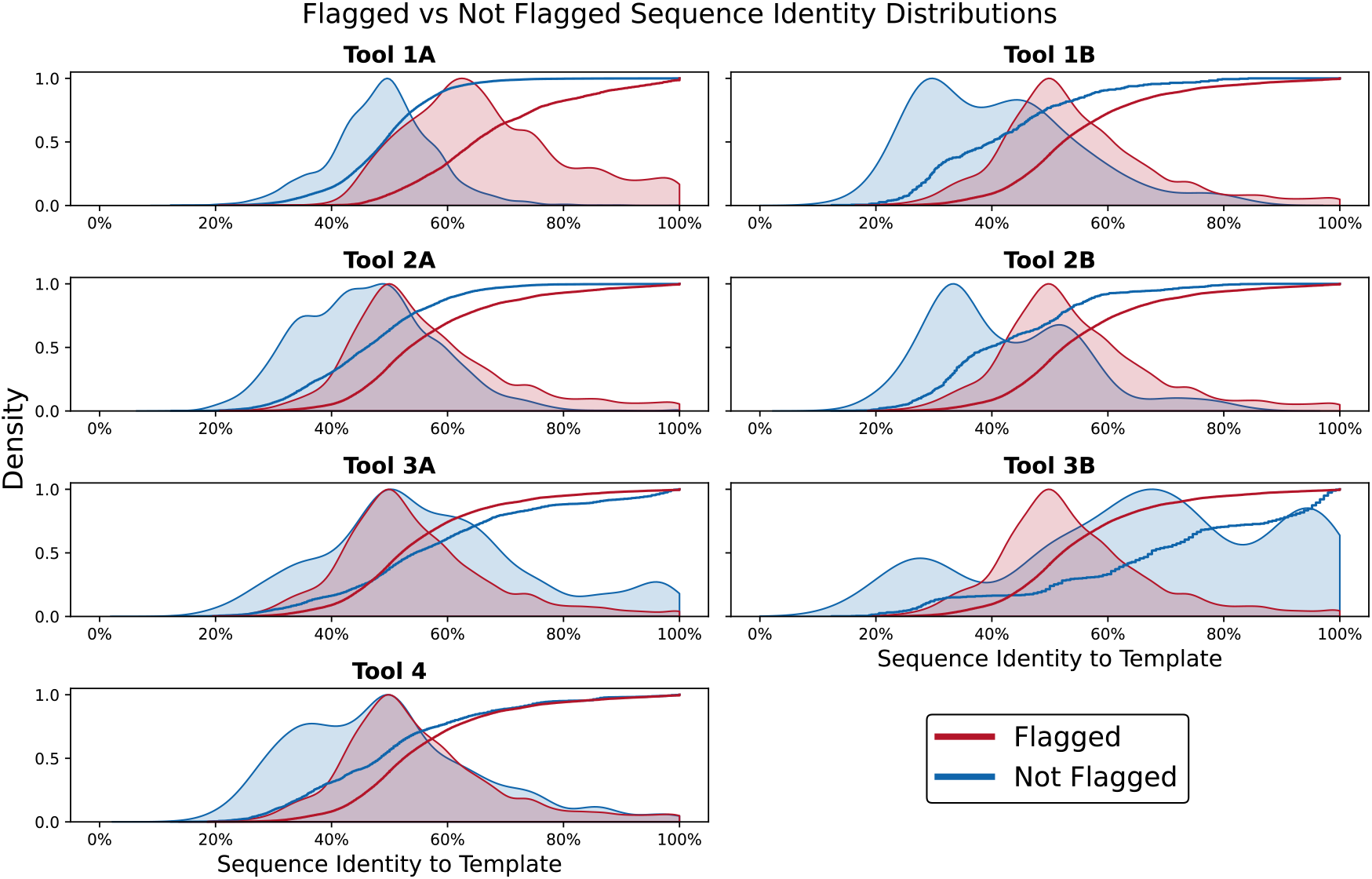
Distribution of sequence identities relative to their templates for reformulated variants with higher probability of maintaining function (defined as having a TM-Score > 0.5 and a ΔpLDDT > -10). Lines in each plot represent an empirical cumulative distribution function (ECDF) of the sequence identities for the variants separated by whether the variants were flagged as potentially hazardous (red) or cleared (blue). Distributions give a kernel density estimate based on the data represented in the ECDFs. Interestingly, while many of the missed variants had low sequence identity relative to their templates, this was not a universal observation, and there was also a sizable contingent of missed proteins with high sequence identity. This is particularly notable for Tools 3A and 3B, where missed variants tended to have higher sequence identity to their templates than flagged ones did.

Notably, the results provide insights into why reformulated variants derived from those templates were difficult to detect. For instance, reformulated variants from Proteins 1 (which were the most-missed variants by Tools 1B, 2B, and 4), 3 (which were frequently missed by Tool 2B), and 4 (also frequently missed by Tool 2B) are derived from proteins of concern with high sequence similarity to proteins that are not of concern. If a reformulated variant is a better match to one of these benign proteins than to a controlled one, then it was not flagged, which is considered the correct outcome for a BSS tool under existing guidance from the U.S. government. As another example, there is disagreement over whether reformulations of Protein 5 (the most-missed variants by Tool 3B and the second-most-missed variants by Tool 4) should be flagged at all, as Protein 5 (1) is only an accessory component of a controlled toxin that provides basic structural integrity with some potential (though nonessential) role in toxin entry and (2) comes from a toxin type that is not typically attributed to human disease. As a final example, variants of Protein 2 (frequently missed by Tool 1B) were not detected as the uniform parameters used for training Tool 1B in this study resulted in poor detection of these proteins. Subsequent adjustment of the parameters for this template indicated that proteins derived from these templates can be detected as well (results not shown).

The identified causes of misses highlight some core learnings from this study: (1) The lack of flagging of synthetic homologs of Proteins 1, 3, 4, and 5 brings attention to the necessity of coming to a consensus on what constitutes a potentially hazardous sequence. Indeed, Provider 4 opted not to update their software precisely because of the lack of clarity regarding whether the proteins “missed” by their tool should have been captured at all. (2) The lack of flagging of variants of Protein 2 shows that, while an approach may generally work well, rather than a one-size-fits-all approach, at least some degree of customization may be needed for detection of specific threat classes.

### Challenge of Obfuscated AI-Reformulated Proteins

So far, all results presented are from BSS runs applied to DNA that *directly* encoded either a protein of concern or a reformulated version of a protein of concern. Unfortunately, we cannot always expect a bad actor to provide nucleic acids that would encode their intentions so clearly. Indeed, effective BSS must also be able to identify nucleic acids that *indirectly* encode a wild-type or reformulated protein of concern, i.e., nucleic acids that could be manipulated using molecular biology techniques to produce a gene that directly encodes a biohazardous protein.^19^

Figure 5 shows the frequency with which the synthetic homologs with high in-silico metrics from each of the template proteins were obscured and revealed by such a DNA obfuscation process for each BSS, where an “obscured” protein is one that was flagged when directly encoded by DNA but not when encoded by obfuscated DNA and a “revealed” protein is the opposite: one that was *not* flagged when directly encoded but that *was* flagged when encoded by obfuscated DNA. Overall, we can see that, along with having a greater ability to detect AI-reformulated variants, all BSS strategies now used by the screening providers have an improved resilience to DNA obfuscation relative to the ones that were in use when the framing study was performed, with the rates of obscuration and revealment dropping for all BSS tools.

**Figure 5.**
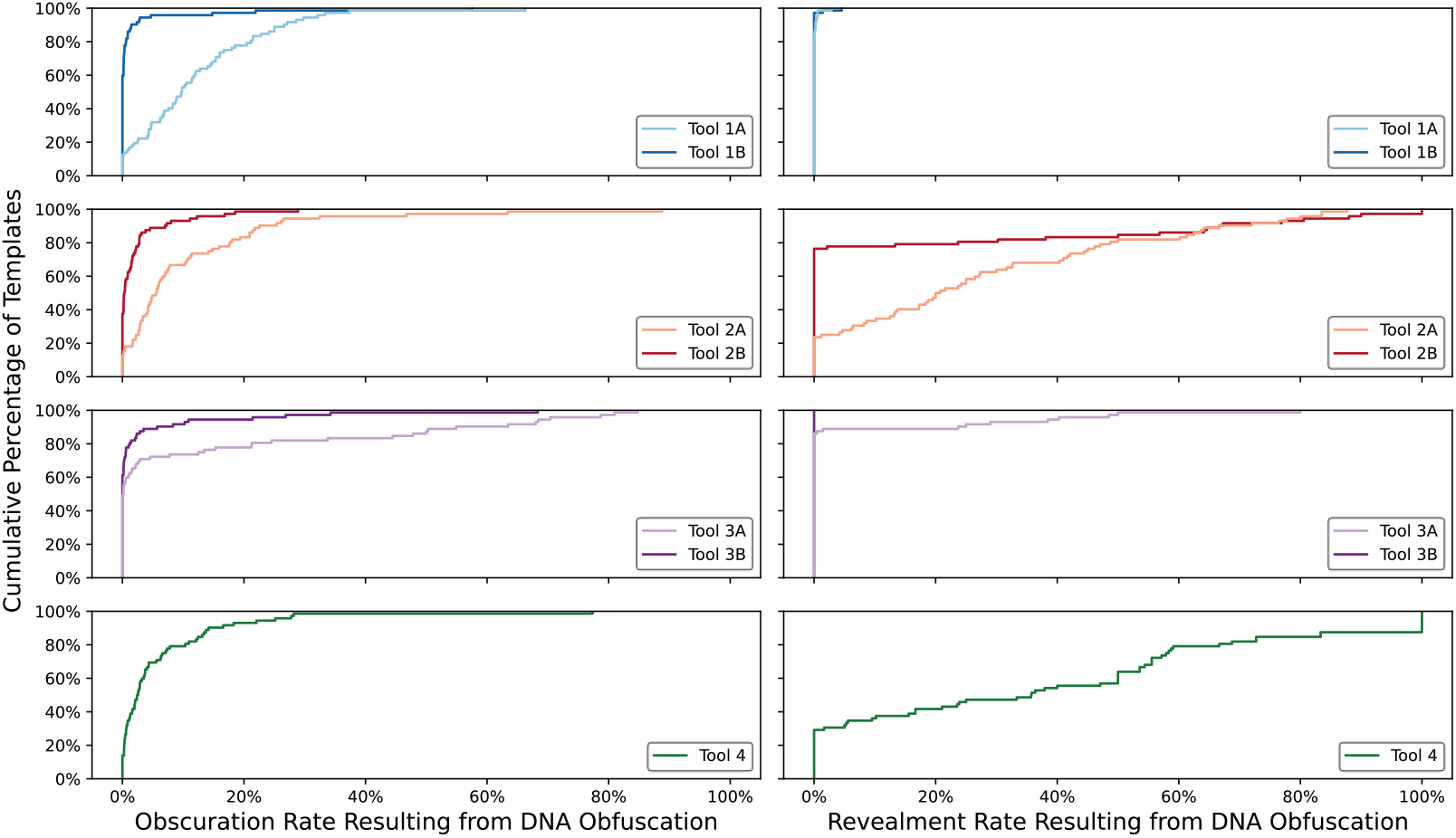
Effects of DNA obfuscation on biosecurity screening software (BSS) for synthetic homologs with high in-silico metrics (TM-Score > 0.5 and ΔpLDDT > -10). Each row gives results for a different BSS provider. Left column: Empirical cumulative distribution functions (ECDFs) of the frequencies with which reformulated proteins were obscured by DNA obfuscation, where an “obscured” protein is one that was flagged when directly encoded by DNA but not when encoded by obfuscated DNA. Right column: ECDFs of the frequencies with which reformulated proteins were revealed by DNA obfuscation, where a “revealed” protein is one that was not flagged when directly encoded but that was flagged when encoded by obfuscated DNA. In both columns, the y-axis indicates the percentage of templates that have a rate lower than the x-axis value, e.g., for Tool 1A, looking at Column 1, approximately 80% of the templates have an obscuration rate of less than 20%. In other words, steeper (more left-shifted) curves indicate better performance.

While it is encouraging to see the rate of obscuration decrease in the updated tools, it is important to realize that, in the ideal case, no proteins are obscured. That is, the ideal ECDFs in the first column of Figure 5 would just be vertical lines at x = 0%, indicating no obscured reformulated proteins regardless of the wild-type template used to generate those proteins. Indeed, by recalculating the confusion matrices from Table 1 using the BSS results from the obfuscated DNA (Table 2), we see that the abilities of all tools to distinguish between the wild-type controlled sequences and protein sequences generated with no conditioning information other than length were minimally affected by DNA obfuscation. This, combined with the fact that the rate of flagging the more-probably-functional sequences decreases for all tools tested, tells us that the combination of AI reformulation and DNA obfuscation is more challenging for BSS than either exploit alone. Still, even with the minor drop in performance that results, the tools perform well, with their “miss” rates against more-probably-functional synthetic homologs remaining comparable to the false negative rates against the wild-type proteins of concern.

**Table 2.** Confusion matrices for each tested biosecurity screening software (BSS) against protein sequences encoded by obfuscated DNA. This table follows the same layout as Table 1.

|  |  |  | More Probably Functional? |  | Control Type |  |
| --- | --- | --- | --- | --- | --- | --- |
|  |  |  | - | + | - | + |
| Label Predicted by BSS | Tool 1A | - | 0.8495 | 0.7133 | 0.9969 | 0.0247 |
|  |  | + | 0.1505 | 0.2867 | 0.0031 | 0.9753 |
|  | Tool 1B | - | 0.5998 | 0.0190 | 0.9792 | 0.0062 |
|  |  | + | 0.4002 | 0.9810 | 0.0208 | 0.9938 |
|  | Tool 2A | - | 0.7078 | 0.2000 | 0.9998 | 0.0247 |
|  |  | + | 0.2922 | 0.8000 | 0.0002 | 0.9753 |
|  | Tool 2B | - | 0.4609 | 0.0334 | 0.9984 | 0.0000 |
|  |  | + | 0.5391 | 0.9666 | 0.0016 | 1.0000 |
|  | Tool 3A | - | 0.6974 | 0.1759 | 0.9984 | 0.0741 |
|  |  | + | 0.3026 | 0.8241 | 0.0016 | 0.9259 |
|  | Tool 3B | - | 0.5945 | 0.0417 | 0.9984 | 0.0185 |
|  |  | + | 0.4055 | 0.9583 | 0.0016 | 0.9815 |
|  | Tool 4 | - | 0.5544 | 0.0938 | 0.9847 | 0.0062 |
|  |  | + | 0.4456 | 0.9062 | 0.0153 | 0.9938 |

## Discussion

To our knowledge, this study represents the first large-scale, coordinated effort between multiple nucleic acid synthesis companies and BSS providers to evaluate new and emerging biosecurity risks introduced by AIPD tools. Unsurprisingly to experts across both the protein engineering and biosecurity fields, our results confirm that AIPD can introduce new potential biosecurity risks, as evidenced by the initial failures in our framing study of two widely used BSS to detect synthetic homologs of a protein of concern and by the varied capabilities of a wider set of BSS to detect synthetic homologs derived from a wider set of potentially biohazardous proteins. That said, we also found that these risks can be mitigated with an AI red-teaming process of adversarial generation of candidates, stress-testing BSS, insight building, and updating or “patching” BSS. Much like the way in which BSS methods adapted when codon engineering became readily accessible, BSS tools can also be modified to be more resilient to the low-sequence-identity synthetic homologs that can be produced by modern AIPD tools.

It is important to highlight some caveats regarding the interpretation of this study. First, as noted throughout this paper, we operate without ground-truth data regarding the functionality of our synthetic homologs, instead relying on in-silico metrics assumed to be correlated to the probability that a given synthetic homolog will maintain function. While literature evidence, personal communication with other AIPD experts, and unpublished results from an analogous follow-on study using benign proteins as proxies for hazardous proteins suggest that these metrics can be useful for classifying non-functional versus potentially functional synthetic homologs, there is no direct confirmation of their effectiveness on the proteins studied here. We do not know how many of our “more-probably-functional” variants would actually be functional in vitro. Still, even if most designs are nonfunctional now, given the rapid pace of advancement in AI and AIPD, it is not difficult to imagine a near future where most designs work. It is far better to have these “AI-resilient” screening mechanisms in place now, when the overall risk is low, than scrambling to make a patch later when the threat is more salient.

It is also important to note that real-life order screening involves both customer screening and sequence screening of an entire order (*i*.*e*., not just one sequence). Because success rates for AI-generated protein sequences remain low, a bad actor would likely need to order numerous sequences before obtaining a functional one. If just one of those sequences were flagged, the actor may be detected and thwarted. Thus, a sequence-by-sequence red teaming exercise like that performed here does not precisely mimic the expected realities of orders and should, in fact, be considered as evaluating one of the most challenging situations in which BSS can be deployed.

In the long run, we should not expect that sequence-based strategies alone—like those tested and developed in this study—will be sufficient, as we envision a future in which AI-assisted generation of proteins produces sequences unlike any seen in nature. Thus, in addition to continuing to improve sequence-based hazard detection software, we must also pursue the development of new approaches. For instance, next steps could include investigating and optimizing the utility of homology detection via learned embedding spaces for augmenting existing BSS.^45–49^ Beyond machinery for screening of nucleic acid orders by BSS, institutional and community strategies will also be important to pursue,^29^ such as improvements to record-keeping of orders that would enable better coordination and follow-up by nucleic acid synthesis companies and other relevant bodies if misuse of synthetic nucleic acids was ever suspected.

We stress that any next steps must be carried out responsibly. Follow-up work exploring biosecurity vulnerabilities and mitigations resultant from advances in AI technologies should coordinate with the appropriate government security authorities in advance of studies. The red teaming of dual use biological capabilities can be a danger in itself, and so such exercises require a formalized assessment of risks and benefits. Red teaming explorations should be designed to minimize risks of studies and experiments. For example, studies involving physical wet-lab experimentation should use benign proxies of threat agents whenever possible and engage in legal reviews that can explore constraints, such as those imposed by export controls.

For the effort presented here, practices adapted from cybersecurity were a starting point and motivated many of the decisions we made regarding how to organize and complete this work. Practices borrowed from cybersecurity include pursuing external red teaming only after exhausting one’s own internal red teaming efforts, conducting new red teaming only after addressing key vulnerabilities that are already known, and not publishing details of vulnerabilities that are difficult to patch. The authors of the framing study involved the nucleic acid synthesis companies and shared concerns and planned experiments *before* running any tests on the BSS used by those companies. Once the results of those tests confirmed the presence of potential vulnerabilities, the leads of the framing study informed the appropriate individuals at relevant regulatory and industry bodies about the potential threat and the planned follow-on studies. Details of the potential vulnerability were kept confidential until patches could be developed, validated, shared, and deployed. In addition to alerting and informing members of the IGSC, members were provided with access to diagnostic test sequences. We are now working with an array of other organizations to provide broader access.

This work shines light on an urgent need to develop community norms around AI biosecurity red teaming, coupled with rapid responses to biosecurity screening vulnerabilities as they are identified.^50^ Biosecurity screening would benefit enormously from development of supporting communities, institutions, standards, training, and practices like those found in the cybersecurity community. National governments have several critical roles to play, including incentivizing adherence to screening best practices, as the U.S. has done recently in the OSTP Framework on for Nucleic Acid Synthesis Screening in allowing scientists to use federal research dollars only at nucleic acid providers that screen appropriately. Governments must also be proactive in funding studies, basic methods research, and ongoing capability assessments to monitor alignment between emerging AIPD capabilities and BSS defenses.

Given the deeply international nature of the nucleic acid synthesis industry, regulatory approaches mandating specific screening requirements are most powerful when they are adopted by the widest array of countries possible. International organizations like the International Biosecurity and Biosafety Initiative in Science (IBBIS) can advocate for the adoption of such uniform regulatory approaches across multiple countries, serve as a neutral third party for reporting suspicious orders, and engage in long-term educational outreach to providers and customers. There is also a role for national and international standards bodies in coordinating the development of metrics and best practices for assuring BSS functionality. The ISO 20688-2 standard, released in March 2024, focused on product quality control, but a section on biosafety and biosecurity requires that “all DNA producers should use a sequence screening mechanism to evaluate ordered sequences” and could serve as a focal point for international standardization of screening.^51^ Both of these types of organizations can also coordinate technical community contributions across national boundaries and help to serve as a long-term, independent repository for technical know-how and guidance on both AIPD misuse risk as well as state-of-the-art screening.

We believe that the ongoing advancement of AIPD holds great promise for addressing some of humanity’s most pressing challenges in the health and life sciences, with the potential to have an overwhelmingly positive impact on people and society.^52^ As with other emerging technologies, however, it is also crucial to continue identifying and mitigating new risks associated with capabilities introduced by harnessing AI methods in the biosciences. We hope this project serves as a timely example of how responsive and responsible risk mitigation can be achieved by combining technical efforts, collaborative exploration, and a thoughtful, deliberative process, thereby contributing to the growing body of work on AI safety.^37^

## Methods

### Variant Generation

Owing to its potential as an information hazard, details regarding variant generation are withheld.

### Screening DNA for Potentially Hazardous Proteins

The four biosecurity screening tools (considering original and patched versions to be the same tool) used in this study are Aclid, BBN’s FAST-NA Scanner, IBBIS’s Common Mechanism, and Battelle’s UltraSeq. We specifically do not identify which results are from which tool in order to focus on the general problem of resilience to AIPD-reformulated variants, rather than a critique or comparison of specific tools.

#### Aclid

Aclid is a security and compliance automation platform for providers of synthetic DNA and RNA. Providers use Aclid to screen and verify customers, screen DNA and RNA for pathogenic and toxic elements, and automate trade compliance. Aclid employs a sequence-based approach to threat detection using sequences and annotations derived from public and proprietary experimental, simulated, and predicted datasets. The screening system includes curated databases of sequences and functional annotations, hardware-accelerated sequence similarity and alignment tools, and algorithms to interpret compliance requirements from sequence data.

Results were generated using Aclid’s cloud-based screening service using the generated sequences in FASTA format as provided without modification. Outputs were downloaded in an open-source, column-oriented data storage format called Apache Parquet and converted to a comma-separated text file. Positive labels were assigned to sequences identified as controlled by Aclid’s screening system and negative labels were assigned to all others.

Aclid’s screening system defines controlled sequences compliant with the definitions in the White House’s Framework for Nucleic Acid Synthesis Screening and the Screening Framework Guidance for Providers and Users of Synthetic Nucleic Acids.^24,31^ In particular, controlled sequences must be a best match to a regulated agent in a 50-nucleotide window.

#### Common Mechanism (IBBIS)

The International Common Mechanism for DNA Synthesis screening uses a combination of profile HMM and BLAST searches to identify potentially hazardous proteins.^53^ While BLAST-based searches are appropriate for identifying proteins with a best taxonomic match to regulated pathogens, HMMs are designed to detect remote functional homologs to proteins of interest, making them well-suited to detecting synthetic homologs.

Profile HMMs were initially only built for proteins from regulated pathogens held in publicly available virulence factor databases and toxins with models in the Pfam database. The detection of the synthetic homologs evaluated in this study varied considerably between proteins with a dedicated HMM and those being detected using BLAST only, with HMMs detecting many more AI-generated sequences.

To examine the uplift in screening accuracy provided by more comprehensive coverage of hazard proteins by HMMs, models were constructed using homologous protein families with high similarity to wild-type sequences for each protein. The expansion of the HMM database resulted in more comprehensive detection of AI-reformulated sequences.

#### FAST-NA Scanner (RTX BBN Technologies)

FAST-NA Scanner uses Bloom filters to identify nucleic acid and amino acid k-mer “signatures” that are unique to specific types of pathogen or toxins, then scans input sequences for these signatures.^54,55^ To make FAST-NA Scanner AI-resilient, signatures were generalized from k-mers to regular expressions.

#### UltraSeq (Battelle)

The core UltraSEQ engine was used as described in Gemler et al., skipping sample genomics preprocessing and downstream metagenomics services.^56^ The standard UltraSEQ databases were used (UniRef100 database,^57^ the SoC database,^58^ a pathogen-specific nucleotide database, and internal ribosome entry site sequences). As described in Gemler et al., UltraSEQ automatically identifies regions within a sequence through the Query Mapper Service and applies a threat score for each region (i.e., subsequence) identified using a logical rules engine.^56^ Specifically for sequence screening, we used UltraSEQ’s Gene Screening Rules Engine as described elsewhere,^59^ flagging anything with a risk level of 5 or below (using default settings of L_min_>50 nts and C_min_>0.2).

#### In-Silico Evaluation of Generated Proteins

All synthetic homologs were evaluated in silico using an open-source protein structure prediction tool. This tool returns (1) predicted 3D-coordinates of each non-hydrogen atom in the protein and (2) a measure of confidence in the returned coordinates for each residue in the form the predicted local distance difference test (pLDDT).

Using predicted structures and pLDDTs, three in-silico metrics were calculated for each synthetic homolog:

1. The structure-level predicted local distance difference test (pLDDT). This is calculated as the mean of the pLDDTs returned for each residue:

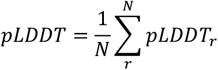

where *N* is the number of residues in the protein. A higher pLDDT is typically reported if the sequence for which a structure is being predicted shares similarities with sequences used to train the structure prediction model. In other words, it can be loosely thought of as a measure of how “realistic” an input sequence is. A higher pLDDT thus indicates a more realistic reformulated sequence and so a sequence more likely to retain function.
2. The predicted template modeling score (TM-Score) between the backbone atoms of the predicted structure of a synthetic homolog and each of 200 predicted structures of its template protein. The TM-Score is a measure of the deviation between coordinates of aligned atoms and is standardized between 0 (exclusive) and 1 (inclusive), where a score of “1” indicates identical structures. The TM-Score between a single synthetic homolog structure and its associated template is calculated as

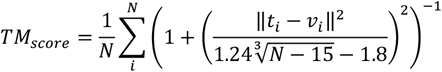

after applying a structural alignment between the template and reformulated structures. In the above equation, *N* is the number of atoms, *t*_*i*_ is the coordinates of backbone atom *i* in the template structure, and *v*_*i*_ is the coordinates of backbone atom *i* in the reformulated structure. A synthetic homolog with a higher TM-Score is predicted to be more likely to share a fold with its wild-type template and so more likely to retain that template’s function. Throughout this paper, the mean of the set of 200 TM-Scores for each reformulated variant is typically reported.
3. The difference in pLDDTs (ΔpLDDTs) resulting from the predicted structure of a synthetic homolog and each of 200 predicted structures of that protein’s template. This is calculated as

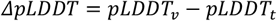

where *pLDDT*_*v*_ is the pLDDT for the synthetic homolog and *pLDDT*_*t*_ is the pLDDT for the template protein. A synthetic homolog with a positive ΔpLDDT can be thought of as being considered more “realistic” and so more likely to maintain function, and vice versa. As with the TM-Score, throughout this paper, the mean of the set of 200 ΔpLDDT scores for each reformulated variant is typically reported.

## Acknowledgments

We thank Ava Amini, David Baker, Charlie Bell, Jonathan Carlson, William Chappell, Oliver Crook, Paul Friedrichs, Steph Guerra, Wendy Hall, Ethan Jackson, Emily LeProust, Sheng Lin-Gibson, Becky Mackelprang, Brad Malin, Jason Matheny, Sarah McGee, Greg McKelvey, Arati Prabhakar, Gopal Sarma, Jonas Sandbrink, Jared Schuetter, Kevin Scott, Lance Stewart, Lynda Stuart, Geoffrey Taghon, Igor Tsyganskiy, Harris Wang, Kevin Yang, and Jaime Yassif, for discussions, guidance, feedback, and assistance. We thank Catherine Romero for her efforts with establishing collaboration agreements among participating organizations. We thank Paul Koch for setting up the secure server used to share sequences, in-silico metrics, and the flags assigned by different tools.

This document does not contain technology or technical data controlled under either U.S. International Traffic in Arms Regulation or U.S. Export Administration Regulations.

## CRediT Taxonomy

BJW: Conceptualization, Data Curation, Formal Analysis, Investigation, Methodology, Project Administration, Resources, Software, Validation, Visualization, Writing – Original Draft, Writing – Review and Editing

TA: Investigation, Resources, Software, Writing – Review and Editing

CB: Data Curation, Formal Analysis, Investigation, Methodology, Project Administration, Resources, Writing – Review and Editing

JB: Formal Analysis, Investigation, Methodology, Project Administration, Resources, Software, Writing – Review and Editing

AC: Formal Analysis, Investigation, Methodology, Software, Writing – Review and Editing

JD: Investigation, Methodology, Project Administration, Writing – Review and Editing

KF: Formal Analysis, Investigation, Methodology, Resources, Software, Writing – Review and Editing

BTG: Formal Analysis, Investigation, Methodology, Resources, Software, Writing – Review and Editing

TM: Formal Analysis, Investigation, Methodology, Resources, Software, Writing – Review and Editing

STM: Formal Analysis, Investigation, Methodology, Resources, Software, Writing – Review and Editing

NW: Formal Analysis, Investigation, Methodology, Resources, Software, Writing – Review and Editing

EH: Conceptualization, Formal Analysis, Investigation, Methodology, Project Administration, Resources, Visualization, Writing – Original Draft, Writing – Review and Editing

## Appendix Framing Study

What follows is the confidential writeup from October 2023 describing the formulation of synthetic homologs and the challenge of detecting them with the DNA screening software employed at the time by two prominent DNA synthesis providers. The methods and results described in the whitepaper served as framing for the confidential process that was undertaken over the following 12 months. Some corrections have been made to the text since its original distribution to fix errors that were subsequently identified. In all cases of correction, the original text has been struck through to indicate a deletion.

## 1 Summary

Over the last five years, artificial intelligence (Al) methods applied to protein design have greatly expanded our ability to engineer proteins. The evolving approaches and trajectory of advancement are providing tools for custom-tailoring proteins to take on numerous challenges in medicine, bioscience, chemistry, and material science. We anticipate that progress in AI-driven protein design will be foundational to breakthroughs in many scientific and engineering fields, with great promise for elevating our quality of life. However, as much as these new Al-assisted protein engineering tools have the power to do good, they can just as easily be used by bad actors to do harm. For instance, using Al methods, it is now possible to dramatically alter a protein’s sequence while preserving its function. We are concerned that this hitherto impossible diversification capability has potential to allow the nefarious generation of variants of dangerous proteins that are undetectable by the hazard detection software used today by synthetic DNA providers to screen customer orders. The new capabilities enhance the ability of would-be bioterrorists to obtain hazardous protein material. We explore a set of methods for performing adversarial Al-assisted protein design (AAIPD) aimed at evading traditional methods for detecting the intention to produce hazardous proteins.

We report on a preliminary investigation into the capabilities of existing hazard detection software used by synthetic DNA providers to identify potentially dangerous proteins after well-known default sequences of amino acids are modified. As an initial AAIPD study, we present several sample methods for producing variants of base proteins and demonstrate in a case study how these methods can be employed to create dissimilar sequences with a high likelihood of retaining the function of dangerous base proteins.

After generating several tens of thousands of alternate sequences of we engaged with Twist Bioscience and Raytheon BBN—two companies that employ nucleotide screening—to understand the value of current methods to identify the modified proteins. We found that existing hazard detection software was incapable of identifying AI-generated variants of, with up to 100% of AI-generated variants of passing through the software undetected. While it is unclear how many of our designed sequences would retain (or, indeed, exceed) native-like function, the methods we use have proven success in other applications, so it is not unreasonable to expect that at least some of the variants that passed the filter would retain toxicity. Regardless, at a minimum, this study indicates the potential of AAIPD to allow potential bad-actors to order genes encoding potentially dangerous variants.

Working with Twist and Raytheon, hazard detection methods were improved to detect hazardous variants produced by AAIPD strategies that employ current AI-assisted protein engineering tools. However, as AI-assisted protein engineering methods continue to improve, hazard detection systems will need to be updated as well. We thus propose an approach to DNA hazard detection analogous to cybersecurity, where white-hat individuals continue to attempt to “hack” their way around existing security measures, then work to address identified vulnerabilities in advance of public disclosure.

## 2 Background

Proteins are macromolecules critical to almost all of life’s processes. The instructions for generating every protein in every organism are encoded in that organism’s DNA. Specifically, in a protein-coding gene, three-letter DNA codons encode one-letter protein amino acids, which are the monomeric units that, when chemically chained together, form a protein. The full sequence of codons in a gene thus defines the full sequence of amino acids in a protein.

To synthesize a protein, biologists and others working with proteins order the DNA sequence that encodes it from one of numerous synthetic DNA providers, then express that synthesized DNA in a host organism. That is, the scientists seeking the protein rely on the biological machinery present in a host organism to build the protein for them using the instructions present in the DNA. To prevent their customers from either accidentally or intentionally ordering DNA that encodes dangerous proteins, synthetic DNA providers have developed software that identifies potentially hazardous sequences and prevents them from being ordered. While the exact strategies used by existing hazard detection software vary from provider to provider, in general, the detection methodology works by looking for similarities between known hazardous DNA sequences (and the proteins they encode) and the ordered DNA.

Recent advances in AI-assisted protein engineering facilitate the diversification of protein sequences associated with known proteins while maintaining function, an engineering capability that was previously extremely difficult. Because existing hazard delection software rely on identifying similarities between known hazardous sequences and ordered sequences, the newfound simplicity with which protein variants can now be designed raises the question as to whether these software are sufficient for threat mitigation. By extension, it also asks how current methods can be extended to be more robust to adversarial manipulations of sequence performed in a manner that does not significantly degrade function.

## 3 Methods

### 3.1 Protein Variant Production

For our case study, we selected as a model can enter the human body by Due to its relatively low LD_50_ and the ease with which it can be distributed is of concern for use in bioterrorism. To build sequence analogs, we

It is also important to note that, because existing DNA hazard detection software rely on calculating similarities between known and potential dangerous sequences, it is reasonable to expect that analogs with less sequence identity to the wild type would be more difficult to detect Thus, there is a direct correlation between the probability that our analogs will maintain function and the probability that they will be detected by hazard detection software: the more residues we fix to the wild-type identity, the greater the sequence identity will be between the variants produced and the wild-type sequence, so although the probability of maintaining wild-type-like function will be improved, the probability of detection by hazard detection software will increase as well.

The expected correlation between detection rates and expected likelihoods of preservation of function is somewhat comforting. We nonetheless have sought a deeper understanding of the degree of diversification of sequences for needed before sequences^ can evade identification by existing hazard detection software. We thus created six sets of ∼ 10k protein variants each, all with differing levels of wild-type residues conserved:

1. Set A:
2. Set B:
3. Set C:
4. Set D:
5. Set E:
6. Set F:

From Set A to Set F, we roughly expect the difficulty of detection to decrease and the probability of function to increase—Set F is thus the one most likely to contain dangerous proteins while Set A is the least likely.

### 3.2 DNA Variant Production

Synthetic DNA providers receive specifications of DNA sequences from their customers, not protein sequences. Thus, the sequences built in Section 3.1 must be reverse translated into DNA before being passed into hazard detection software. This provides another opportunity to test the robustness of these software, as explained in the three methods we used for the reverse translation process:

1. The first strategy produced a baseline: As multiple codons equivalently encode individual amino acids, we replaced the codon for each amino acid in each protein sequence by a randomly selected codon encoding that amino acid.
2. Because codons are 3 letters long, there are 3 possible reading frames along a single-stranded DNA molecule. For instance, the DNA strand ATGGCAAA can be read as ATGIGCA, TGGICAA, or GGCIAAA, corresponding to the three different amino acid sequences MA, WQ, and GK, respectively. Thus, to translate DNA into protein, the reading frame must be known; otherwise, the incorrect protein sequence will be read. Recognizing the above, in the second strategy we prepended a randomly generated DNA sequence for sperm whale myoglobin and a segment of junk DNA sequence for sperm whale myoglobin and a segment of junk DNA to DNA sequences generated using the first approach. Junk DNA was also appended to the DNA sequences
3. Of course, our second reverse translation strategy could be bypassed easily if the hazard detection software simply tested all DNA reading frames rather than (a strategy that is already employed in hazard detection software). To make detection of full analogs impossible in any reading frames, in our last reverse translation strategy,

DNA sequences were prepared for each set using the first method. Additionally, DNA was produced for Set A using methods 2 and 3. The DNA for Set A produced using reverse translation approach 1 was labeled as Set A_0; the DNA produced using reverse translation approach 2 was labeled as Set A_l, and so on.

### 3.3 Submitting DNA to Twist Biosciences and Raytheon BBN

Set A_0 was submitted to Twist Biosciences; all sets of DNA were submitted to Raytheon BBN (the organization that provides DNA hazard detection software to IDT). These two companies use different hazard detection software, though the specific details of how these software function are unknown to us.

## 4 Results

A summary of results can be found in Table 1.

### 4.1 Twist Bioscience

Twist Bioscience submitted DNA Set A_0 to their ThreatSeq software. Of the ∼10k sequences in this set, ThreatSeq detected only one of them as a threat—the wild-type sequence that was included as a positive control. The remaining sequences were not detected as a threat, indicating that, absent any other intervention, they could currently be ordered through Twist Bioscience.

As a follow-up, in the context of our submitted challenge, Twist Bioscience passed all protein sequences of Set A through ESMFold to predict their three-dimensional structures, then aligned those predicted structures against the AlphaFold-predicted structure for to calculate a TM score for each, which is a measure of structural similarity with values on the range (0, 1], where 1 indicates a perfect match between structures. Generally, TM scores below 0.17 correspond to unrelated, randomly chosen proteins while scores above 0.5 correspond to proteins that have the same overall fold. The team found that all of the variant sequences had a TM score above 0.17 and ∼9k had a TM score above 0.5, indicating that ESMFold-predicted structures could identify analogs reliably where the ThreatSeq software could not.

### 4.2 Raytheon BBN

Raytheon BBN submitted all DNA sets to their internal threat detection software. As with ThreatSeq, only the positive control sequence was identified as a threat—all variants went undetected and so could currently be ordered through IDT absent any other intervention.

Following the failure to detect variants, the Raytheon BBN team had an insight regarding another approach that could be more resilient to AI-diversified harmful proteins. The details of the new methods are unknown to us; however, the updated approach was far more effective at identifying the submitted DNA as encoding analogs, as summarized in Table 1. By report of the Raytheon BBN team, the revised threat detection methodology had a false-positive rate (defined as how frequently the system incorrectly identified a random sequence from a set of ∼8k as a analog) of 0.000%.

**Table 1:** Reported detection rate of analogs using the original methods employed by Twist and Raytheon BBN as well as using the new method employed by Raytheon BBN.

| Set Name | Sequence Identity<br>(%, $\mu \pm \epsilon$) | N Variants | Detection Rate<br>Twist (%) | Detection Rate<br>Raytheon Old (%) | Detection Rate<br>Raytheon New (%) |
| --- | --- | --- | --- | --- | --- |
| A_0 | [REDACTED] | 9985 | 0.00 | 0.00 | 100.00 |
| A_1 | [REDACTED] | 9985 | N/A | 0.00 | 100.00 |
| A_2 | [REDACTED] | 9985 | N/A | 0.00 | 99.94 |
| B | [REDACTED] | 10000 | N/A | 0.00 | 100.00 |
| C | [REDACTED] | 10000 | N/A | 0.01 | 100.00 |
| D | [REDACTED] | 10000 | N/A | 0.00 | 100.00 |
| E | [REDACTED] | 10000 | N/A | 0.18 | 100.00 |
| F | [REDACTED] | 10000 | N/A | 4.36 | 100.00 |

## 5 Conclusions and Future Directions

The existing DNA hazard detection systems of both Twist Biosciences and IDT—two of the leading synthetic DNA providers—were unable to identify any AI-diversified analogs. This result clearly demonstrates that, given newfound Al-assisted protein engineering strategies, DNA hazard detection systems that had been in place are insufficient for mitigating bioterrorism threats. We believe the results make salient the need to make changes in our existing threat detection systems to address this shortfall.

Future threat detection methods must satisfy three criteria: (1) the false negative rate must be low (i.e., threats must be detected), (2) the false positive rate must be low (i.e., proteins that are not threats must not be detected), and (3) the method must be computationally inexpensive to operate.

These criteria make the structure-based method proposed by the Twist team a challenge to implement in practice, as predicting protein structure is an expensive computational operation to perform. Not to mention, the third reverse translation system that we describe above could easily bypass a structurebased search. As for Raytheon BBN’s alternative strategy, we do not know the details of it and cannot comment at this time on its broader feasibility. However, we assume that it still relies on a sequence-based search.

If the current rapid pace of advancement in Al-assisted protein engineering continues as is, the nature of the threats with which DNA hazard detection software will have to contend will likely evolve quickly and become ever-more sophisticated. We should not expect that sequence-based strategies alone will be sufficient in the long-term, as we envision a future where Al-assisted generation of proteins produces sequences unlike any seen in nature. In addition to improving sequence-based hazard detection software, then, we should pursue the development of new approaches as well. For instance, there is likely value in investigating homology detection via the learned embedding space of large protein language models, research into which is already thoroughly underway. Similarly, the detection of hazardous proteins among a sea of nonhazardous proteins is at its heart a form of anomaly detection, a problem that has already received extensive attention among the broader machine learning community.

Regardless of the threat mitigation strategies developed to date and in response to the challenges presented in this paper, it is important to recognize that the future of biosecurity will be ever changing. In the same way that cybersecurity teams must constantly develop new strategies to mitigate new threats as new Al approaches are developed, those dedicated to preventing acts of bioterrorism will have to continue to adapt as well.

## 6 Acknowledgments

We thank James Diggans and Emily Leproust at Twist Bioscience, Mark Behlke, Adam Clore, Gavin Kurgan, and Jeff Manthey at IDT, and Jake Beal and Tom Mitchell at Raytheon BBN for participating in this study. In particular, we thank James Diggans at Twist Bioscience and Jake Beal and Tom Mitchell at Raytheon BBN for running the variant sets that we sent them through their respective DNA hazard detection software and analyzing the outputs. We thank David Baker at the Institute for Protein Design at the University of Washington for feedback on design of variant sets.

